# Whole-genome sequencing revealed local genetic differences and multi-layered genetic structure in Asian brown bear (*Ursus arctos*) populations

**DOI:** 10.1101/2025.04.21.649633

**Authors:** Yu Endo, Naoki Osada, Tsutomu Mano, Alexei V. Abramov, Ryuichi Masuda

## Abstract

A multi-layered genetic structure shaped by multiple demographic events across different time periods has been well documented in some species. However, evaluating this pattern remains challenging in many wide-ranging wildlife because genome-wide datasets with broad geographic coverage are still scarce. Brown bears are genetically differentiated among regional populations, but the genomic data are still limited and the demographic events remain incompletely understood. Here, we generated whole-genome resequencing data from brown bears in Asian local populations previously reported to show significant variation and combined these data with ancient samples for population genomic analyses. Our results support previously reported demographic events, such as the early divergence of Central Asian populations. Furthermore, we found genetic differences among Asian brown bear populations, suggesting that population-specific demographic histories occurring at different times have contributed to their unique genetic characteristics. In the Central and West Asian populations, divergence times estimated from autosomal genomes differed by hundreds of thousands of years from those inferred from mitochondrial DNA and Y-chromosomal DNA haplotypes, suggesting that gene flow continued after regional dispersal. In addition, the Y-chromosomal phylogeny and *f*_4_ statistics indicated genetic differentiation between the Kazakhstan and Tibetan populations, likely resulting from male-biased introgression into the Tibetan population. Finaly, the Hokkaido and Etorofu (Iturup) populations retained alleles derived from hybridization with polar bears, likely due to the limited impact of post-LGM population expansion. These findings indicate that the genetic variation among brown bear populations is shaped by multiple demographic events occurring at different times, resulting in a multi-layered genetic structure.

**Significance Statement:** Multi-layered genetic structure is well known in some species, but remains poorly understood in wild species due to limited genomic data. In this study, we newly sequenced nine brown bears in Asia which included bears that genome data in their area is unreported. By combining these with modern and ancient genomes, we conducted population genomic analyses to characterize local genetic variation and detect hybridization with related species such as the polar bear. We identified (1) substantial divergence time gaps in Central and West Asian populations between autosomal genomes and mitochondrial and Y-chromosomal haplotypes, (2) male-biased introgression into the Tibetan population, and (3) limited impact of recent expansion between island and Eurasian continental populations. These findings suggest that the genetic structure of the populations has been shaped by multiple demographic events at different times, resulting in a multi-layered genetic structure and providing new insights into how wildlife genetic diversity is shaped.

## Introduction

Multi-layered genetic structure is an idea that modern genetic characteristic reflected multiple demographic events in different times. This has been described in humans, where the genome has been shaped by multiple migrations and demographic changes though time (e.g., Tassi et al. 2015). For example, genome-wide genetic data revealed that the current Japanese population consists of two main genetic layers from Jomon hunter-gatherers and rice-farming Yayoi migrants, and more genetic layers from different migration events (Osada & Kawai 2021). Although the genetic structure of other species, such as the black rat, *Rattus rattus* (Puckett et al. 2020), the spruce, *Picea abies* and *P. obovata* (Zhou et al. 2024), and the Japanese wolf, *Canis lupus hodophilax* (Segawa et al. 2022), was also proposed to be multi-layered and reflect population demography at different times, the presence of a multi-layered genetic structure in wild species that distribute widely remains largely unknown because there is a lack of global genome-wide data and their limited distribution.

The brown bear, *Ursus arctos,* is a large mammal in Carnivora which distributes widely on the Northern Hemisphere and well known their global scale genetic structure and hybridization with related species. At the inter-species level, they have hybridized with different ursid species such as the cave bear, *Ursus spelaeus* (Barlow et al. 2018) and the polar bear, *Ursus maritimus* (Edwards et al. 2011). In particular, hybridization between polar bears and North American brown bears has been reported in many previous studies. Mitochondrial DNA (mtDNA)-based phylogenetic trees place polar bears within a brown bear lineage (clade 2) that is nowadays distributed in the Admiralty, Baranof, and Chichagof (ABC) Islands (Leonard et al. 2000). In contrast, phylogenetic trees estimated from nuclear genomes clearly showed their differentiation between two species (Hailer et al. 2012; Miller et al. 2012), although alleles derived from the hybridization has been confirmed in brown bear genomes (Cahill et al. 2013, 2015, 2018), and their geographic distribution pattern remains unclear. The direction of introgression between these species remains an interesting topic and has been discussed in several previous studies (Cahill et al. 2013, 2015, 2018; Lan et al. 2022; Liu et al. 2014; Miller et al. 2012; Wang et al. 2022), so we should consider this potential influence when interpreting the results of hybridization with polar bears.

At the intra-species levels, variation among populations is known. Multiple mtDNA lineages with different estimated divergence times have been reported to exhibit distinct geographic distributions (Davison et al. 2011; Hirata et al. 2013; Lan et al. 2017; Figure 1). The appearance and divergence times of lineages have been consistently supported by many previous studies, including those using ancient DNA from different areas (da Silva Coelho et al. 2023; Edwards et al. 2011; Kosintsev et al. 2022; Molodtseva et al. 2022; Segawa et al., 2021). On the other hand, nuclear genomic analysis did not exhibit a clear differentiation between those mtDNA lineages (Bidon et al. 2014; Hirata et al. 2017; Endo et al. 2021; de Jong et al. 2023). de Jong et al. (2023) revealed (1) the autosomal genome reflects recent population demography, and suggested (2) the mtDNA and Y-chromosomal DNA haplotypes distribution was reflected both the older structure or incomplete lineage sorting. These two points cannot be explained by simple relationships in which a single demographic event uniformly shapes genetic variation across populations. Rather, they reflect complex processes, where both the occurrence of historical events and the magnitude of their genetic impacts differ among populations, giving rise to a multi-layered genetic structure. In fact, de Jong et al. (2025) reported that genetic distances among North American brown bear populations reflect both past and recent demographic events. Therefore, greater attention to local populations is necessary to better understand the genetic variation and demographic history of brown bears.

**Figure 1.**
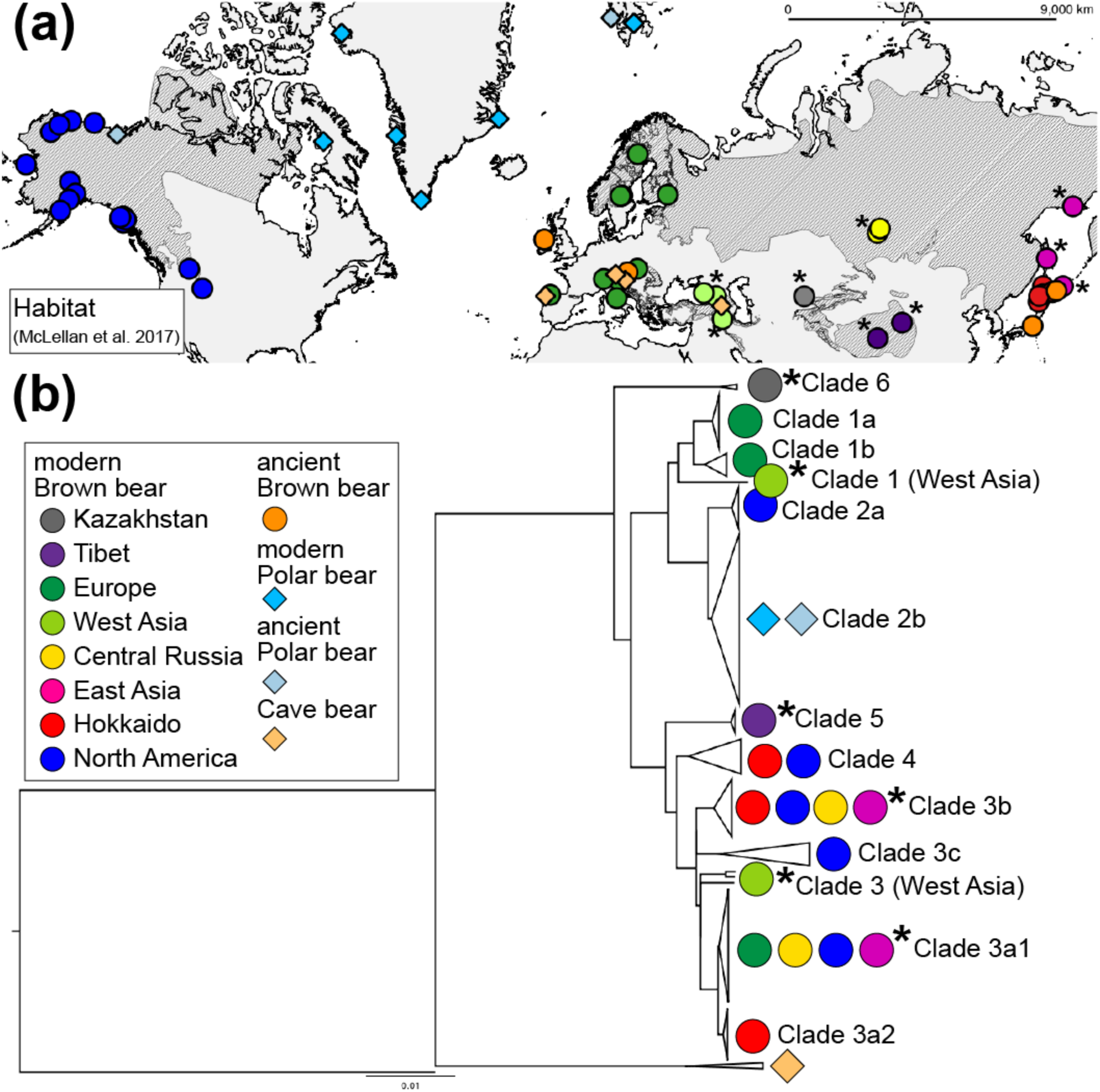
Sampling locations and mitochondrial DNA (mtDNA) lineages found in each location. Sampling locations for bears analyzed in this study excluding outgroups (a) and a phylogenetic tree of brown bear mtDNA lineages (b). The shaded area reflects the current habitat of the brown bear (McLellan et al. 2017). The dot colors represent correspond to the groups listed in Table S1: ancient brown bear, dark orange; ancient polar bear, light blue; cave bear, light orange; brown bears from West Asia, light green; brown bears from Central Asia, purple; brown bears from Central Russia, yellow; brown bears from East Asia, pink; brown bears from Europe, green; brown bears from Hokkaido, red; brown bears from North America, blue. Shapes represent species: square, polar bear or cave bear; circle, brown bear. The asterisks shown near the dots indicate that new sequence data were reported in this article. These sampling locations were inferred based on the collected region (Hirata et al. 2014) and presence of brown bear habitat (McLellan et al. 2017). Two samples (sample name: GT_1 and Asiatic_black_bear) are not shown in this map because these individuals were from zoos.

In this study, we focus on local populations in Asia. These populations exhibit distinct genetic characteristics from those on the Eurasian and North American continents based on mtDNA analyses (Hirata et al. 2013, 2014; Lan et al. 2017; Tumendemberel et al. 2019) and a few individuals exhibit nuclear genomic distinctiveness relative to other populations (de Jong et al. 2023; Tumendemberel et al. 2023). However, genomic data for these populations remain limited. To clarify the variation in local Asian populations and the demographic history underlying this pattern, we performed whole-genome resequencing on nine brown bears, whose mitochondrial DNA lineages previously reported by Hirata et al. (2013, 2014) from the Asian populations (Etorofu (Iturup) Island; Sakhalin; Northern Okhotsk; Kazakhstan; Tibet; Central Russia; and West Asia; Figure 1 and Supplementary table 1). To infer local differences in phylogeny, demography, and gene flow, we conducted population genomic analysis using publicly available modern genomic data (Barlow et al. 2018; Benazzo et al. 2017; Cahill et al. 2013, 2015; Endo et al. 2021; Kumar et al. 2017; Liu et al. 2014; Miller et al. 2012; Taylor et al. 2018; Wang et al. 2022) and ancient genomic data (Barlow et al. 2018; Cahill et al. 2018; Lan et al. 2022; Segawa et al. 2021; Wang et al. 2022).

## Results

We genotyped 42,906,277 SNPs for modern samples and 10,537,259 SNPs for ancient samples. We only considered transversion SNPs for ancient samples to remove deamination mutation. We used two datasets, one including only modern samples (n = 61, 42,906,277 SNPs) and the other including both modern and ancient samples (n = 72, 10,537,259 SNPs). The coverage, which is the proportion of covered bases with depth ≧ 1 to the total bases of the reference sequence (Supplementary table 1), was 10.14%–91.59% for both ancient and modern samples, but 81.26%–91.59% for only modern samples. The depth for modern samples was 6.85–55.11 (Supplementary table 1).

### Genetic diversity, genetic structure, phylogeny, and demography

We computed heterozygosity and runs of homozygosity (ROH) for each individual to assess the genetic diversity (Supplementary figure 1). Bears from Central Asia exhibited high heterozygosity and low ROH, whereas the sample from Etorofu (Iturup) Island displayed the opposite pattern (Supplementary figure 1), a trend that is often observed in small populations.

Principal component analysis (PCA) showed genetic affinity between individuals from geographically close locations (Supplementary figures 2a and 3). The first principal component reflected the genetic difference between the North American individuals and other individuals, whereas the second principal comportment divided the East Asian individuals from the European and West and Central Asian individuals, this was supported by the result of *K* = 4 in a maximum likelihood-based analysis of individual ancestries, ADMIXTURE (Supplementary figure 4). In addition, the third and fourth principal components showed local distinction of the Kazakhstan and Tibetan individuals from other individuals (Supplementary figure 2b), and this was also shown in PCA based on the modern dataset (Supplementary figure 3) and *K* = 3, 4, 7, and 8 in ADMIXTURE (Supplementary figure 4). This result is consistent with previous studies (de Jong et al. 2023; Tumendemberel et al. 2023), which reported that individuals from or near Kazakhstan and Tibet are deeply diverged from other populations. The relative effective migration rate around Asia region especially central to east Asia was estimated as lower than other areas by Estimated Effective Migration Surfaces, a method that infers spatial variation in effective migration rates based on genetic data (EEMS; Supplementary figure 4). Isolation of the Hokkaido, East Asia, and West Asia populations was also supported by the result of *K* = 2 in ADMIXTURE (Supplementary figure 4), where the cross-validation error was the lowest (Supplementary figure 4g). The results of *K* = 3 and 5 showed two Central Russian individuals differed in their genetic affinities to other groups (Supplementary Figure 4b and d).

Phylogenetic trees based on mtDNA, X-chromosomal, autosomal, and Y-chromosomal DNA were constructed to investigate the identity of differentiation between the populations (Figure 2). Compared with trees based on the mitochondrial DNA showing the geographic discontinuities of mtDNA haplotypes (Davison et al. 2011; Figure 1B), the trees based on the nuclear genome reflected affinity between nearby populations (Figure 2); this was also, as previously reported by de Jong et al. (2023). Individuals from Kazakhstan and Western Asia, who had localized mtDNA haplotypes not observed elsewhere, tended to be distinct from other individuals, such as those from Europe, North America, and other Asian areas. Individuals from Tibet also exhibited this trend. However, only Y-chromosomal phylogenetic tree showed that they were included within the same lineages as the European, North American, Central Russia, and East Asian individuals (Figure 2d). The mtDNA lineages found in West Asia (clade 1_West Asia and clade 3_West Asia) were different from other lineages reported in previous studies (e.g. Davison et al. 2011), and both diverged earlier than each related lineage (clade 1 and clade 3, respectively). All phylogenies showed that the two Central Russian individuals diverged at different times. Their divergence times of clade 1_West Asia were 253 thousand years ago (95%CI: 143–374 thousand years ago; Supplementary figure 5). That of clade 3_West Asia 139–146 thousand years ago (95%CI: 73.7–220 thousand years ago; Supplementary figure 5), which approximately corresponds to the divergence time of the Y-chromosomal West Asian lineage, estimated at 158 thousand years ago (95% CI: 140–177 thousand years ago; Supplementary figure 6). In contrast, the divergence time based on autosomal genome between Central Asia (CAsia_Tibet_U05M, CAsia_Kazakhstan_U13M, CAsia_Tibet_U40) and West Asia (WAsia_U12M, WAsia_U47M, WAsia_Ge), estimated using SMC++, was substantially more recent than those inferred from the mtDNA and Y-chromosomal DNA haplotypes (64,577 years ago; Table 1) although it was earliest among the five groups pairs. In addition, SMC++ estimated that Western Asia and Europe diverged 21,099 years ago (95%CI: 15,485-23,758 years ago; Table 1). The Hokkaido population (Hokkaido_421, Hokkaido_518, Hokkaido_819) was estimated to have diverged from the North American population (NorthAmerica_BB020, NorthAmerica_BB037, NorthAmerica_GP01) before the Last Glacial Maximum (LGM; 16,187 years ago; 95% CI: 13,307–20,293 years ago; Table 1) whereas divergence from continental East Asia (EAsia_AA1, EAsia_U34) may have occurred after the LGM (7,410 years ago; 95% CI: 2,332–13,396 years ago; Table 1).

**Figure 2.**
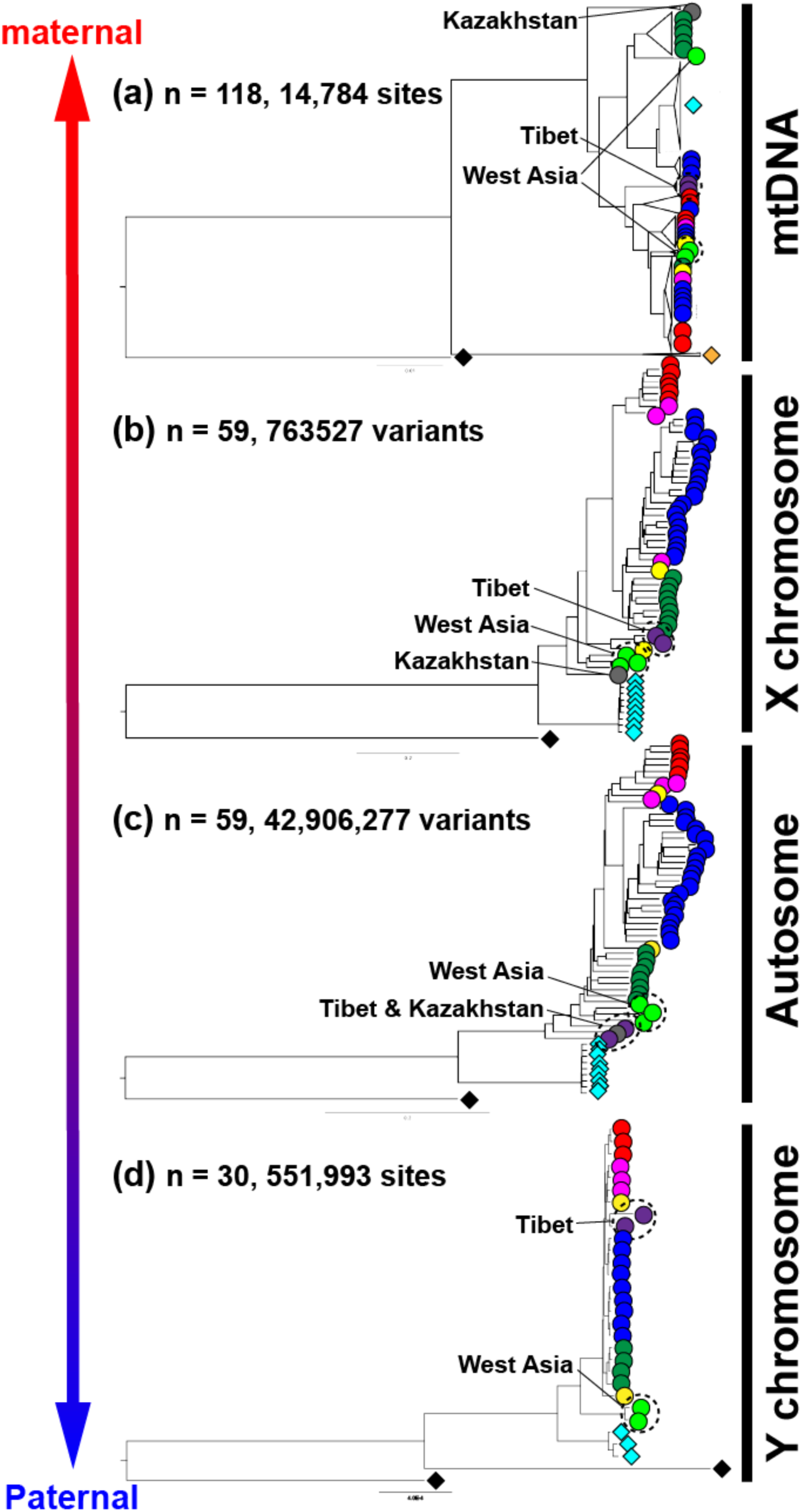
Phylogenetic trees for four different heritable regions: mtDNA (a), the X chromosome (b), autosomes (c), and the Y chromosome (d). The number of individuals and total sites or variants are shown in each tree. The dot shape and color correspond to those in Figure 1. Colors represent the groups listed in Table 1: ancient brown bear, dark orange; ancient polar bear, light blue; cave bear, light orange; brown bears from West Asia, light green; brown bears from Central Asia, purple; brown bears from Central Russia, yellow; brown bears from East Asia, pink; brown bears from Europe, green; brown bears from Hokkaido, red; brown bears from North America, blue. Shapes represent species: square, polar bear or cave bear; circle, brown bear.

Pairwise sequentially Markovian coalescent analysis (PSMC) was performed on modern brown bear samples to estimate their population size (Supplementary figure 7). Changes in effective population size were similar among individuals from Europe, North America, and Kazakhstan (Europe_SLK1, NorthAmerica_ABC1, and CAsia_Kazakhstan_U13M), whereas the Tibetan individual (CAsia_Tibet_U05M) was notably different from others (Supplementary figure 7a, b). Nearly identical trends were observed between the Hokkaido and Etorofu (Iturup) Island individuals (Supplementary figure 7c), and between the West Asian and European individuals (Supplementary figure 7d). The population size changes in the Northern Okhotsk, Sakhalin, and Hokkaido were similar until approximately 100 thousand years ago; subsequently, but thereafter, the effective population sizes in the Northern Okhotsk and Sakhalin were slightly larger than that in Hokkaido (Supplementary figure 7e).

### Inter- and intra-species gene flow

Population splits and mixtures were inferred using TreeMix (Supplementary figure 8). The phylogeny optimally estimated based on Δ*m* (Supplementary figure 9) revealed a genetic affinity between nearby populations, which was consistent with the findings from the phylogenetic tree (Figure 2). The results of migration edges = 1, 2 indicated that migration occurred from the ABC island population to the polar bear population. Additionally, the result of migration edges = 5 showed the migration from the polar bear population to the Hokkaido and Etorofu (Iturup) Island populations.

To detect hybridization with related species, *f*_4_(Outgroup, Polar bear; X, Kazakhstan[CAsia_Kazakhstan_U13M]) and *f*_4_(Outgroup, Cave bear; X, modern Polar bear), were calculated, where X represents brown bear individuals. Assuming that the target related species is equally diverged from X and Kazakhstan[CAsia_Kazakhstan_U13M] or modern Polar bear, and that there is no gene flow between the target related species and Kazakhstan[CAsia_Kazakhstan_U13M] or modern Polar bear, using American black bear and Asiatic black bear as an outgroup. Almost values of *f*_4_(Outgroup, Polar bear; X, Kazakhstan[CAsia_Kazakhstan_U13M]) were significantly negative excluding *f*_4_(Outgroup, Polar bear; West Asia, Kazakhstan[CAsia_Kazakhstan_U13M]) (Supplementary table 3, |Z score| > 2). It was also supported when using the only modern samples dataset (Supplementary table 4). The result of *f*_4_(Outgroup, Cave bear; X, modern Polar bear) show that statistical significance varied depending on the outgroup used. Only *f*_4_(American black bear, *kudarensis* Cave bear; Kazakhstan[CAsia_Kazakhstan_U13M], modern Polar bear) and *f*_4_(American black bear, *kudarensis* Cave bear; West Asia, modern Polar bear) were statistically significant (|Z score| > 3; Supplementary table 5), although *f*_4_(Asiatic black bear, *kudarensis* Cave bear; Kazakhstan[CAsia_Kazakhstan_U13M], modern Polar bear) and *f*_4_(Asiatic black bear, *kudarensis* Cave bear; West Asia, modern Polar bear) were not.

To compare the difference between populations regarding hybridization with related species, we calculated *f*_4_ ratio and found a geographically discontinuous pattern in the proportion of hybridization with polar bears. In instances where X represented an individual with a statistically significant *f*_4_ value, the proportions of gene flow estimated by *f*_4_ ratio (Figure 3; Supplementary figure 10; Supplementary table 6, 7) from polar bears or cave bears to brown bears were at most 24.36% and 13.81%, respectively. Although the *f*4 ratio strictly assumes one-directional gene flow with nothing else going on, so the values of *f*4 ratio could be overestimates if gene flow of the opposite direction was also present, the *f*_4_ ratio (Figure 3), showed results consistent with the *f*_4_ statistics (Supplementary table 3 and 4). Specifically, it indicated that (1) hybridization was geographically discontinuous from North America to Hokkaido and Etorofu (Iturup) Islands, (2) ancient Japanese brown bears showed a higher proportion of loci from hybridization than modern Japanese brown bears, and (3) these patterns remained unchanged when ancient polar bear genotype data were used instead of modern polar bear data.

**Figure 3.**
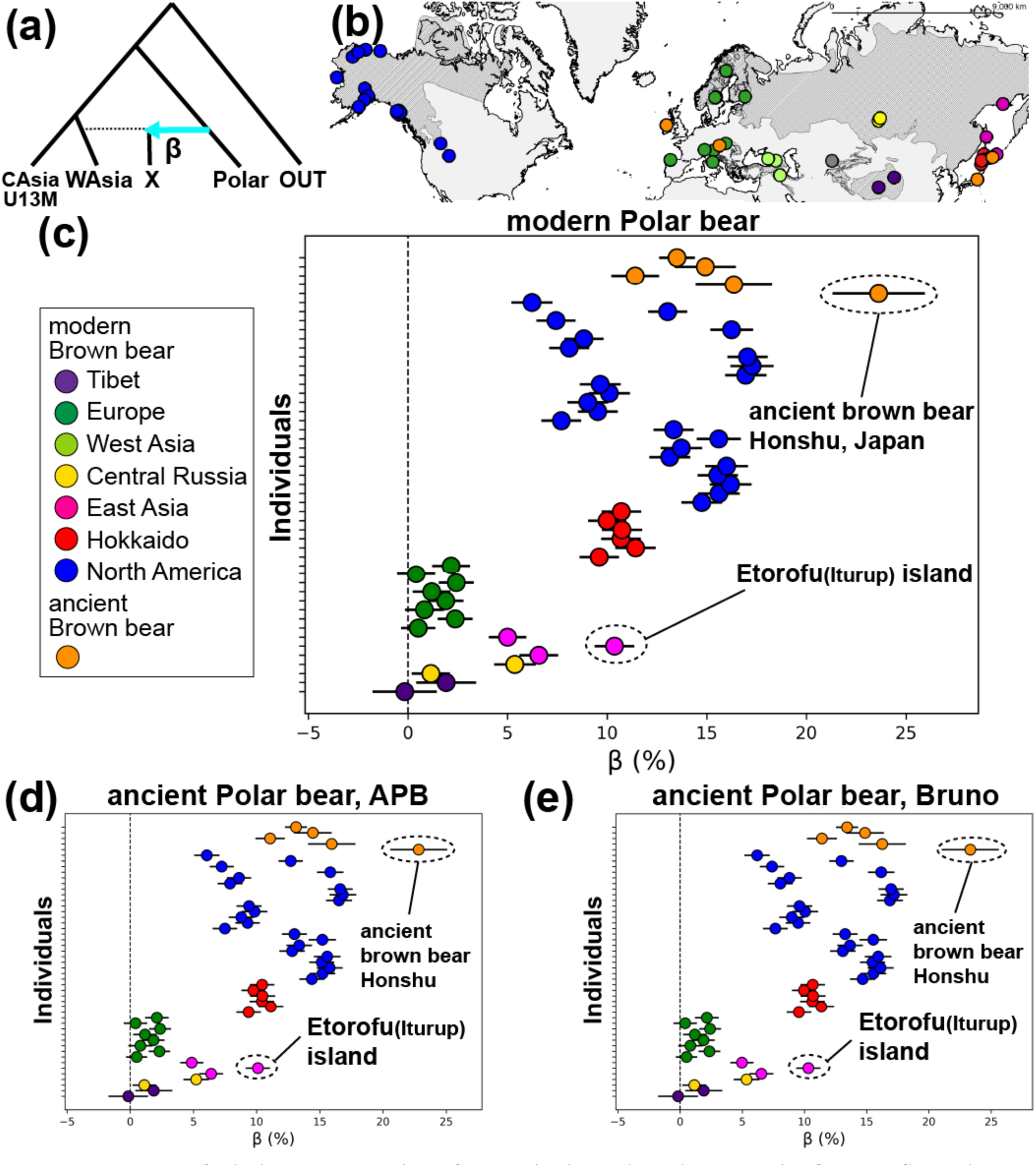
Percent of admixture proportions from polar bears based on *f*_4_ ratio. β (%) reflects the proportion of gene flow from the polar bear to X, representing a brown bear individual (a). The dot colors correlate with those on the world map (b). Colors represent the groups listed in Table 1: ancient brown bear, dark orange; ancient polar bear, light blue; cave bear, light orange; brown bears from West Asia, light green; brown bears from Central Asia, purple; brown bears from Central Russia, yellow; brown bears from East Asia, pink; brown bears from Europe, green; brown bears from Hokkaido, red; brown bears from North America, blue. Each figure shows the result when the polar bear is the modern polar bear (c), APB is an ancient polar bear (d), and Bruno is an ancient polar bear (e). The name of the ancient Polar bear corresponded to the sample name listed in Table 1. The shaded area in Figure 3b reflects the current habitat of the brown bear (McLellan et al. 2017). The tree topology shown in Figure 3a follows that presented to Figure 2 of Patterson et al. (2012).

The timing of hybridization between the polar bear and three brown bears from Hokkaido (Hokkaido_421), Etorofu (Iturup) Island (EAsia_ETF1), and North America (NorthAmerica_GP01) was estimated using MSMC-IM. The ancestors of all three brown bear lineages were inferred to have hybridized with the polar bear approximately 100–200 thousand years ago after speciation (Supplementary Figure 11), although this estimation does not completely correspond with a previous study (Wang et al. 2022).

We calculated *f*_4_(Outgroup, Central or West Asia; East Asia, other groups) to assess differences in gene flow between the Tibetan population and other Central and West Asian populations. *f*_4_(Outgroup, Tibetan individual; East Asia, other groups) values were significantly negative regardless of the outgroup or comparison group used (Supplementary table 8).

## Discussion

In this study, we conducted whole-genome sequencing of nine Asian brown bears to investigate variation and the underlying demographic history of local populations across Asia. Our results support previous findings, including the earlier divergence of Central Asian populations and the retention of alleles derived from hybridization with polar bears in the Hokkaido population. In addition, we identified further demographic events contributing to genetic variation in Asia. Notably, we observed (1) substantial divergence time gaps in Central and West Asian populations between autosomal genomes and mtDNA and Y-chromosomal DNA haplotypes, (2) male-biased introgression into the Tibetan population, as indicated by the Y-chromosomal DNA phylogeny, and (3) a limited impact of recent population expansion with no complete isolation between island and Eurasian continental populations, resulting in the retention of alleles derived from hybridization with polar bears. Overall, our findings suggest that the genetic variation among Asian brown bear populations cannot be explained by a single demographic event. Rather, it reflects multiple events that occurred at different times and together resulted in a multi-layered genetic structure (Figure 4).

**Figure 4.**
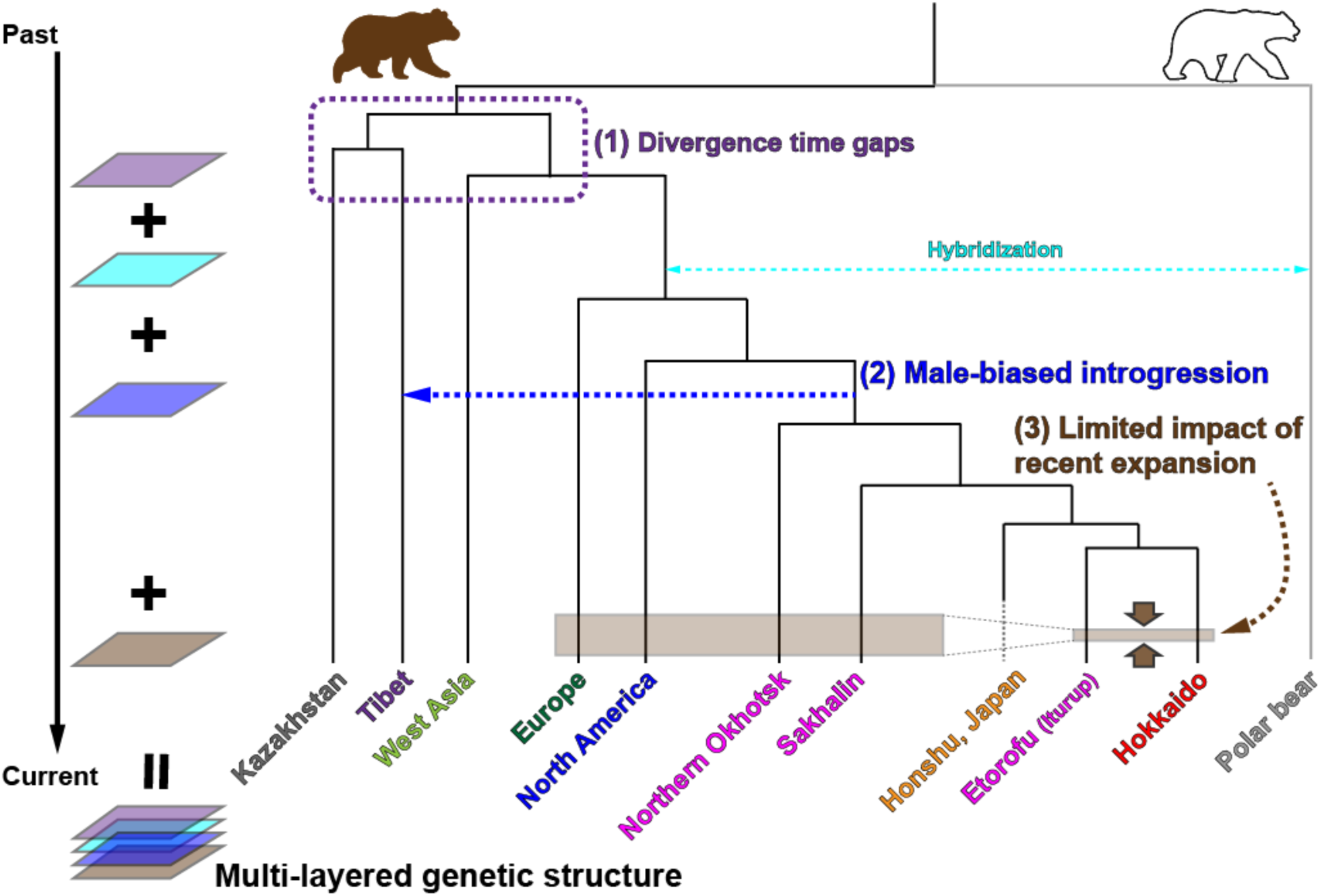
Schematic of genetic variation among Asian brown bear populations. This study indicated (1) substantial divergence time gaps in Central and West Asian populations between autosomal genomes and mitochondrial and Y-chromosomal lineages (purple line), (2) male-biased introgression into the Tibetan population (blue arrow), and (3) limited impact of recent expansion between island and Eurasian continental populations (brown shading), with shading width representing the magnitude of the impact.

### Genetic diversity of the brown bear

Genetic diversity in brown bears from Western and Central Asia was relatively higher compared with that in brown bears from other areas. Genetic structure analysis using EEMS showed that the low migration around Western and Central Asia restricted gene flow from others populations and contributed to the significant distinctiveness of these populations, especially in Kazakhstan, Tibet, and Western Asia; however, the population size changes around Western and Central Asia was inferred to have been constant, which retained genetic diversity. In contrast, Tumendemberel et al. (2023) reported low heterozygosity in brown bears from Mongolia and Pakistan, located relatively close to Kazakhstan and Tibet, and suggested the need for conservation of these populations. Even with high genetic diversity, comprehensive monitoring research is needed in Western and Central Asia to protect this populations from extinction.

In contrast to the high genetic diversity of West and Central Asian brown bears, that of the Etorofu (Iturup) Island individual was significantly lower than that of the Hokkaido individuals and as low as the Apennine brown bear, which was recently listed as endangered; however genetic affinity was also found with the Hokkaido population and this result was consistent with a prior mtDNA analysis (Hirata et al. 2013). These results indicate that the Etorofu (Iturup) Island population experienced a bottleneck after isolation from the Hokkaido population. Events such as the founder effect and other disturbances, including anthropogenic impacts (Kostenko et al. 2004) could be causes of the low genetic diversity. There are approximately 60–260 brown bears on each island in the Kuril Archipelago (Kostenko et al. 2004). Therefore, conservation plans and additional genomic data are needed to monitor this population and evaluate its genetic diversity.

### Demographic history of Western and Central Asian populations

PCA clearly distinguished the Tibet and Kazakhstan individuals from the other populations, and this was also reflected in the phylogenetic tree. Previous studies (Hirata et al. 2013; Lan et al. 2017) reported localized mtDNA haplotypes not observed elsewhere: clade 5 in Tibet and clade 6 around Himalayas. Ancestral genetic traits from those mtDNA lineages have remained in this area because of geographical isolation from other populations, and this inference is partially supported by the result of our ADMIXTURE analysis and a previous study (de Jong et al. 2023).

Comparison of these populations revealed clear genetic differentiation between the Kazakhstan and Tibetan individuals, as indicated by the PCA. The Y-chromosomal DNA phylogenetic analysis indicated that the Tibetan brown bears diverged more recently than the West Asian ones, whose divergence timing was similar to that of the Kazakhstan bear based on autosome phylogeny. In addition, the value of *f*_4_(Outgroup, Central or West Asia; East Asia, other groups) provided significant evidence of gene flow between the East Asian populations and the Tibetan individuals. This pattern suggests that the Tibetan population was more affected by recent male-biased migration than the ancestors of the Kazakhstan individual. Thus, the genetic differentiation of the Tibetan brown bears may be attributable to differences in their demographic history. The PSMC results also supported this interpretation, showing clear differences in effective population size changes between populations. However, the sequencing depth of the Tibetan individual (CAsia_Tibet_U05M) was relatively low (Supplementary Table 1), which may have led to an underestimation of heterozygosity and consequently biased the population size estimates. Further analyses are warranted to further elucidate these trends because the current sampling was limited.

PCA, ADMIXTURE, PSMC, and TreeMix indicated that the Western Asian population was relatively similar to the European populations. However, the phylogeny based on Y-chromosomal DNA sequences revealed that the Western Asian individuals diverged earliest among the brown bear lineages examined in this study (Figure 2) with an estimated divergence time of 158 thousand years ago (95% CI: 140–177 thousand years ago; Supplementary figure 6). The localized mtDNA haplogroups, which were also reported in previous studies (Ashrafzadeh et al. 2016; Hirata et al. 2014; Mizumachi et al. 2020) was estimated that they diverged approximately 100–200 thousands year ago (Supplementary figure 5), a timeframe closer to that of the Y-chromosome haplogroup. Consequently, we infer that the Western Asian population diverged from the European population before the recent expansion and has remained isolated from other populations. However, the divergence time between Western Asia and Europe (21,099 years ago; Table 1) suggesting that limited gene flow may have persisted after divergence, and that the populations were not completely isolated.

The brown bear populations in Central Russia (Krasnojarsk Province, located in Siberia) were proposed to have resulted from admixture between other populations (Europe, North America, and Hokkaido populations). We found that the individuals from this area exhibited high genetic diversity (Supplementary Figure 1), and that the two individuals differed in both their genetic affinities to other groups (Supplementary Figure 4b and d) and their divergence timing (Figure 2). Previous study (Hirata et al. 2014) showed that both major mtDNA lineages, clade 3a1 and clade 3b, with clade 3b diverging earlier than clade 3a1, were found around this area. In a nearby region, northern Mongolia, Tumendemberel et al. (2023) reported a mtDNA lineage that diverged earlier than clade 3, although there were low bootstrap values. Based on these finding, the populations around Central Russia likely formed by admixture between ancestral and recently expanded populations.

*f*_4_ and *f*_4_ ratios using the American black bear as the outgroup detected hybridization between *kudarensis* cave bear, and the brown bear in Western Asia and Kazakhstan. The cave bear is widely distributed on the Eurasian continent several hundred thousand years ago (Knapp et al. 2009); therefore, hybridization could have occurred during that period. Barlow et al. (2021) proposed that there was gene flow between cave bears excluding *kudarensis* cave bear and the common ancestor of the European and Western Asian populations. Our results may reflect additional gene flow from cave bears. However, *f*_4_ value using the Asiatic black bears as the outgroup were not statistically significant. Gene flow between Asiatic black bears, American black bears, brown bears, and polar bears was previously estimated by *D* statistics (Kumar et al. 2017; Barlow et al. 2018; Puckett et al. 2025). This gene flow could be a cause of the statistical significance of *f*_4_(American black bear, Cave bear; X, modern Polar bear) because the value of *f*_4_(W, X; Y, Z) statistics also become significantly negative if there is gene flow between W and Z. Although the statistical significance of *f*_4_ values when other bear species were set as outgroups remains a question, alleles from hybridization with cave bears might have persisted, especially in these populations.

### Demographic history of Etorofu (Iturup) Island and Hokkaido

TreeMix (migration edges = 5) and *f*_4_ statistics confirmed more hybridization between polar bears and brown bears in Hokkaido and Etorofu (Iturup) Islands than in other Eastern Asian areas (Sakhalin and the Northern Okhotsk), which is partially consistent with the results of de Jong et al. (2023), who also showed gene flow between the Hokkaido population and polar bears. Previous studies (Cahill et al. 2013, 2015) identified that, among brown bear populations, the North American population, particularly individuals around the ABC Islands, harbored the most alleles from hybridization with the polar bear. These finding indicate geographically discontinuous gene flow between the polar bears.

Two scenarios regarding this discontinuity are considered: (1) admixture between the Eurasian continental population and a recently expanded brown bear population that was not involved in admixture with polar bears as suggested by de Jong et al. (2023); and (2) additional gene flow between polar bears to only the Hokkaido and Etorofu (Iturup) Island populations. In this study, scenario (1) appears to be more likely based on three results. First, the PSMC results showed that the effective population size changes of the mainland individuals (Sakhalin and northern Okhotsk) have been slightly larger since approximately 100,000 years ago, whereas that of the Hokkaido and Etorofu (Iturup) Island populations has remained constant. It suggests that the mainland ancestors experienced the effects of recent expansion, whereas the ancestors of the Hokkaido and Etorofu (Iturup) Island populations were affected to a much lesser extent. This pattern suggests that the mainland ancestors experienced the effects of recent population expansion, whereas the ancestors of the Hokkaido and Etorofu (Iturup) Island populations were affected to a much lesser extent. Although there is a time gap between the inferred demographic change and the timing of the recent expansion, the discrepancy may be explained by the limited resolution of PSMC for recent demographic events (in humans, approximately within the last ∼30,000 years; Schiffels and Durbin 2014). Second, ancient brown bears tended to share more alleles from hybridization with the polar bear than modern brown bears distributed near where they were collected, and this was, also supported by Cahill et al. (2018). Thus, it is more likely that the brown bear ancestor possessed more alleles from the hybridization with polar bears than modern brown bears, and the recent expansion replaced those alleles with the alleles observed today. Finally, MSMC-IM analysis estimated that the ancestors of brown bears from Hokkaido, Etorofu (Iturup) Island, and North America hybridized with polar bears approximately 100–200 thousand years ago following speciation (Supplementary Figure 11). Although the peak migration rate was not identical between the Etorofu Island individual and the other individuals, the observed difference was not significant. Analyses of additional Etorofu individuals are needed to determine whether further hybridization occurred. This study concluded that the greater retention of alleles derived from hybridization in the Hokkaido and Etorofu (Iturup) Island populations may be attributable to the weaker impact of recent population expansion.

However, this does not imply that the modern Hokkaido and Etorofu (Iturup) Island populations have completely escaped the effects of recent expansion. Based on the *f*_4_ ratio estimates, the ancient brown bear from Honshu, Japan, exhibited a higher proportion of hybridization than the modern Hokkaido and Etorofu (Iturup) Island populations. In addition, the Hokkaido population was inferred to have diverged from continental East Asia relatively recently (7,410 years ago; Table 1), which is slightly later that the disappearance of the Soya land bridge between Hokkaido and Sakhalin (approximately 12,000 years ago; Ohshima 1990), suggesting that limited migration may have occurred in the recent past.

### Multi-layered genetic structure could explain the variation of the brown bear

This study revealed genetic variation and underlying demographic histories among regional brown bear populations, which cannot be explained by a single demographic event. For example, divergence times estimated from autosomal genomes differed significantly from those based on mitochondrial DNA and Y-chromosome haplotypes, suggesting continued gene flow after regional dispersal. The Tibetan population showed slight genetic differentiation from Kazakhstan individuals despite a similar divergence time, likely reflecting male-biased gene flow after divergence, as inferred from discordance between autosomal and Y-chromosomal phylogenies. In addition, post-LGM expansion was restricted between island and Eurasian continental populations, resulting in the persistence of alleles derived from hybridization with polar bears in the Hokkaido and Etorofu (Iturup) populations.

Based on these findings, we propose multi-layered genetic structure of the brown bear genome (Figure 4); it includes both the retention of ancestral genetic traits and replacement of traits with new ones because of the recent expansion is, reflected in the contemporary genetic structure, is plausible to explain the variation among brown bear’s regional population. The reason for the difference in influence of the recent expansion could have primarily arisen because of climate changes. Climate change produces sea-level fluctuations that can lead to land bridges and glacial ice covering land. These environmental changes facilitated migration to new environments while occasionally imposing restrictions. The diverse environmental factors in each area underscore the significance of focusing on local demographic history to more fully elucidate brown bear evolution.

Although our study revealed the variation in brown bear populations, it is important to note that not all brown bear populations were included. For example, individuals from the northern part of Russia, Kunashiri Island, and around the border between East Russia and China were not included because of a lack of samples. Such sampling biases, referred to as Wallacean shortfall, have the potential to overlook genetic diversity and pose challenges in assessing future extinction risks (Lomolino et al. 2016). Additionally, this study and other studies (Cahill et al. 2018; Segawa et al. 2024) showed genetic differentiation between modern and ancient populations, and provided valuable insights into the evolution of the brown bear. Integrating genomic data from individuals across various distribution areas and time periods would offer a more comprehensive understanding of brown bear evolutionary processes and contribute to the conservation of their genetic diversity.

## Materials and methods

### Sampling and whole-genome resequencing

Nine skin or muscle samples were used that were collected by the Zoological Institute, Russian Academy of Sciences (ZIN, Saint Petersburg, Russia; Figure 1 and Supplementary table 1) from the local populations distributed in the Eurasian continent and islands: Etorofu (Iturup) Island, Sakhalin, Northern Okhotsk region, Kazakhstan, Tibet, Central Russia, and West Asia, across brown bear habitat (McLellan et al. 2017). From the samples, we extracted the total DNA using the DNeasy Blood & Tissue Kit (Qiagen, Hilden, Germany) and stored it at 4°C or −20°C until used.

Then, 150-bp pair-end whole-genome resequencing of these samples was conducted using an Illumina Novaseq 6000 sequencer by Macrogen Japan (Tokyo, Japan). The libraries were prepared using the TruSeq DNA Nano Sample Prep Kit (Illumina, San Diego, USA). The average of the total amount of sequence data per sample was ordered 90 Gb.

### Mapping and genotyping

Whole-genome sequence data from the nine brown bears and 63 previously sequenced bears (Miller et al. 2012; Cahill et al. 2013, 2015, 2018; Benazzo et al. 2017; Kumar et al. 2017; Barlow et al. 2018; Taylor et al. 2018; Endo et al. 2021; Segawa et al. 2021; Lan et al. 2022; Wang et al. 2022) were checked for sequence quality, and poor-quality sequences were removed using FastQC ver. 0.11.8 (Babraham Bioinformatics 2011) and Trimmomatic ver.0.39 (Bolger et al. 2014). We mapped raw sequence data to the reference sequence of the polar bear (GCF_017311325.1) using the BWA-MEM algorithm implemented in the BWA ver. 0.7.17 (Li & Durbin 2009) with the ‘-M ‘command option.

To determine sex for individuals (excluding ancient samples specified in Supplementary table 1), we calculated the depth ratio on the scaffolds of the X chromosome (n = 5; NW_024423319.1–NW_024423323.1) and the Y chromosome (n = 19; NW_024423324.1– NW_024423342.1) using the following formula. If the depth ratio was > 1, we set the sex as female.

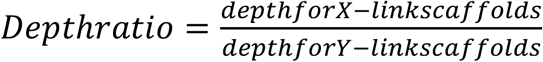

After mapping, single nucleotide variants of modern samples (n = 61, Supplementary table 1) were called and then hard filtered under the following parameters: significant Fisher strand test (FS) > 60.0; variant confidence/quality by depth (QD) < 2.0; RMS mapping quality (MQ) < 40.0; strand odds ratio (SOR) > 9.0; MQRankSum < 12.5, and significant read position bias (ReadPosRankSum) < 8.0 in GATK ver. 4. 1. 2. 0 (McKenna et al. 2010). For single nucleotide variants on the X chromosome scaffolds, genotyping of males was performed with the option --ploidy 1, and that of females with the option --ploidy 2 in GATK. Genmap ver. 1.3.0 (Pockrandt et al. 2020) was used to calculate genome mappability and exclude variants with (30, 2)-mappability < 1.

A different method was used to genotype ancient samples because they had lower sequence quality compared with modern samples. Ancient samples (n = 11; Supplementary table 1) were genotyped by mpileup in samtools ver. 1.9 using the parameters -q 30 -Q 30 and pileupCaller (https://github.com/stschiff/sequenceTools) with the parameters --randomHaploid --skipTransitions after mapping to the reference sequence. Such samples are sometimes not suitable for population genetic analysis because of the small amount of sequence data, so we calculated the ratio of the number of covered bases with depth ≥ 1 to the total bases of the reference sequence using a samtools ver. 1.15 command, samtools coverage. We retained samples with a ratio of > 0.1.

EIGENSOFT ver. 7.2.1 (Patterson et al. 2006), which was used to perform PCA, does not accept data that contain more than 100 chromosomes. Therefore, for *f*_4_ statistics and ancient DNA analysis, we used only scaffolds longer than 0.1 M bp and excluded the sex-linked scaffolds (n = 79) whose total length was approximately 99% of the total reference sequence length because most scaffolds were too short to be used for population genetic analysis.

### Genetic diversity, genetic structure, phylogeny, and demography

We calculated heterozygosity and ROH per individual using PLINK ver. 1.9 (Purcell et al. 2007), using the parameters --het for heterozygosity, and --homozyg-kb 500 and --homozyg-kb 2000 for ROH. The genetic structure of the brown bears (n = 54) worldwide was estimated by PCA using EIGENSOFT ver. 7.2.1 (Patterson et al. 2006) and ADMIXTURE ver. 1.3.0 (Alexander et al. 2009). SNPs with high linkage disequilibrium (*R*^2^ > 0.5 within a 50-SNPs window) and those with minor allele counts < 2 were removed using the following options (--indep-pairwise 50 5 0.5 --mac 2) prior to running PCA and ADMIXTURE. We selected 20 individuals (Europe_191Y, Europe_OFS01, Europe_RF01, WAsia_Ge, WAsia_U12M, WAsia_U47M, Rus_235, Rus_U76, CAsia_Kazakhstan_U13M, CAsia_Tibet_U05M, CAsia_Tibet_U40, EAsia_U34, EAsia_UAA1, EAsia_ETF1, Hokkaido_421, Hokkaido_518, Hokkaido_819, NorthAmerica_ABC01, NorthAmerica_BB020, NorthAmerica_BB037) for ADMIXTURE because it is sensitive to number of samples per cluster. ADMIXTURE was run on each of 2–8 clusters (*K*). For each *K*, 10 independent runs were performed under the default convergence criterion, in which the log-likelihood increased by less than 10^−4^ between iterations (-C 0.0001). The run with the lowest cross-validation error was selected as the final result. The relative scale of gene flow based on genetic distance and sampling location was calculated by EEMS (Petkova et al. 2016). The number of modeled demes was set to be 400 and Markov chain Monte Carlo (MCMC) chains were each run for 20 million iterations; the first 10 million generations were discarded as burn-in and sampling occurred every 10,000th generation.

To construct phylogenetic trees based on autosomal and X-chromosomal SNPs, the identity-by-state of each pair was calculated using PLINK with the parameter (--distance square 1-ibs). We interpreted the non-sex-linked scaffolds as autosomal scaffolds (n = 3,875). After calculating the genetic distance matrix, phylogenetic trees were constructed with the neighbor-joining method (Saitou & Nei 1987) using the ape package in R (Paradis et al. 2004). We used the Asiatic black bear sample (Asiatic_black_bear; Kumar et al. 2017) as an outgroup.

PSMC (Li & Durbin 2011) was performed on new 7 samples (CAsia_Tibet_U05M, CAsia_Kazakhstan_U13M, EAsia_AA1, EAsia_ETF1, EAsia_U34, WAsia_U12M, WAsia_U47M) and 5 samples from the same or near sampling locations (Europe_SLK1, Hokkaido_421, Hokkaido_518, NorthAmerica_ABC1, Rus_235). The methodology was basically following by Endo et al. (2021). Input consensus sequences were generated BCFtools ver. 1.19 (Danecek et al. 2021) mpileup with the option (-C 50). PSMC was run with 25 iterations, a maximum 2*N*_0_ coalescent time of 15, an initial theta/rho ratio of 5, and 64-time intervals using the following parameters (-N25 -t15 -r5 -b -p "4+25*2+4+6"). Bootstrap was performed with 100 replications for each sample. The mutation rate and generation time were set to 1.82*10^-8^ per site per generation and 11 years, respectively (Benazzo et al. 2017; Endo et al. 2021).

Divergence times between groups (CAsia-WAsia, WAsia-Europe, Europe-NorthAmerica, NorthAmerica-Hokkaido, Hokkaido-EAsia) were estimated using SMC++ (Terhorst et al. 2017). We selected variants with a minor allele count greater than 1 and a missing rate lower than 0.2 per scaffold (>1 Mb) using vcftools with the following options (--mac 1 --max-missing 0.8). Input files were generated for five groups (CAsia: U05M, U13M, U40; WAsia: U12M, U47M Ge; Europe: OFS01, RF01, 191Y; NorthAmerica: BB020, BB037, GP01; Hokkaido: 421, 518, 819; EAsia: AA1, U34) and SMC++ was run with the mutation rate and generation time set to 1.82 * 10⁻⁸ per site per generation and 11 years (Benazzo et al. 2017; Endo et al. 2021), respectively, consistent with those used in the PSMC analysis. Bootstrap was performed with 100 replications, and bootstrap input files were generated using the online script bootstrap_smcpp.py (https://github.com/popgenmethods/smcpp/files/1182555/bootstrap_smcpp.zip).

### Mitochondrial DNA and Y chromosomal DNA phylogeny

To obtain the whole mtDNA sequence, whole-genome sequence data from 37 individuals (Supplementary table 1) was assembled using getOrganelle ver. 1.7.7.0 (Jin et al., 2020). The script get_organelle_from_reads.py was run with a maximum of 10 extension rounds (-R 10) and the target organelle genome type set to animal mitochondrial DNA (animal_mt). The k-mer sizes were set to 21, 45, 65, 85, and 127 (-k 21,45,65,85,127). Then, 81 samples (Supplementary table 2) from the NCBI database were downloaded and added as sequence data. We extracted sequence data from two rRNA genes, 22 tRNA genes, and 12 coding genes, but we did not include NADH dehydrogenase subunit 6 sequences because of its nucleotide composition heterogeneity (Hirata et al. 2013), and gap sites were excluded. The final sequence length was 14,784 bp.

The phylogenetic tree was constructed by the maximum likelihood method using iqtree ver. 1.6.12 with the parameters (-m MFP -AICc -bb 1000 -alrt 1000) after alignment with the MUSCLE algorithm (Edgar 2004). The Asiatic black bear sample (Honshu; NCBI accession number: DRR250459) was used as an outgroup.

The divergence time for each mtDNA lineage was then estimated using BEAST ver. 2.4.3 (Bouckaert et al. 2014) under the GTR substitution model. The relaxed lognormal clock and the coalescent constant population model were used. To calibrate divergence times, radiocarbon dates for ancient sequences were set, which is a mean age of 120 thousands of years ago (kya) for the ancient polar bear (GU573488; Lindqvist et al. 2010), and 31.87 thousand years ago, 44 thousand years ago, and 409 thousand years ago for the cave bears (EU327344, NC011112, and KF437625; Bon et al. 2008, Krause et al. 2008, Dabney et al. 2013). Trees were sampled every 10,000 generations from a total of 1,000,000,000 generations; the first 100,000,000 generations were described as burn-in. The maximum credibility tree was generated using TreeAnnotator in BEAST. Other options were followed by the “Divergence Time Estimation” tutorial by Taming the BEAST (Barido-Sottani et al. 2018).

The Y-chromosome sequences were constructed from a .vcf file using bcftools ver. 1.9 (Danecek & McCarthy 2017). We extracted coding regions, and the final sequence length was 551,993 bp. Phylogenetic trees based on haplotype data were constructed following the same method as that of mtDNA using iqtree. The divergence time was estimated using BEAST under a mutation rate of 1.3*10^−9^ per site per year (Wang et al. 2022; de Jong et al. 2023). The other parameters followed the same method as that of mtDNA.

### Inter- and intra-species gene flow

TreeMix ver. 1.13 (Pickrell & Pritchard 2012) was used to show the phylogenetic relationships and gene flow between groups or individuals. After running TreeMix, we inferred the most likely number of migration events using OptM package in R (Fitak 2021).

Hybridization with related species and gene flow between brown bear populations were evaluated by *f*_4_ statistics using Admixtools (Patterson et al. 2012). *f*_4_(Outgroup, Polar bear; X, Kazakhstan[U13M]) and *f*_4_(Outgroup, Cave bear; X, modern Polar bear), where X represents the ancient and modern brown bear individuals, were calculated to detect hybridization with polar bears and cave bears, respectively. An alternative evaluation to determine hybridization with related species, *f*_4_ ratio (Patterson et al. 2012), was used to infer the mixing proportions of an admixture event. We calculated *f*_4_ values, α and β, for each related species as shown below.

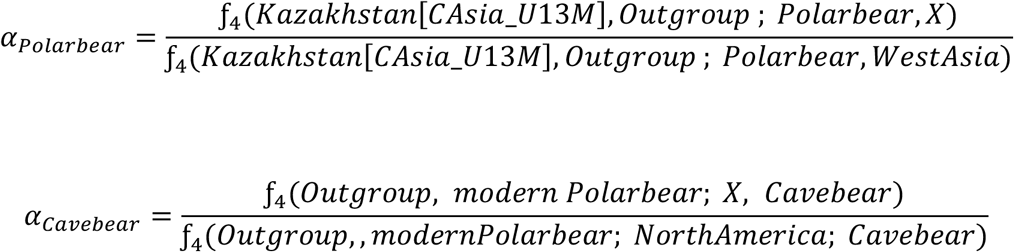

The percentage of the admixture proportion was then calculated following the below formula.

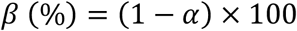

The timing of hybridization with the polar bear was estimated using MSMC-IM (Wang et al. 2020) to examine differences in timing among brown bear groups. We selected one polar bear (PB10) and three brown bears from Hokkaido (Hokkaido_421), Etorofu (Iturup) Island (EAsia_ETF1), and North America (NorthAmerica_GP01). Variants were called using BCFtools mpileup with the options (-q 20 -Q 20 -C 50) and the bamCaller.py script from the MSMC toolkit (https://github.com/stschiff/msmc-tools). Only variants on scaffolds longer than 1 Mb, consistent with those used in SMC++, and with (30, 2)-mappability = 1 were retained. Genotypes were phased using WhatsHap ver. 2.8 (Patterson et al. 2015) with default settings. Coalescence rates between the polar bear and each of the three brown bear populations were estimated using MSMC2, based on MSMC (Schiffels and Durbin 2014), with the mutation rate and generation time set to the same values as those used in PSMC. MSMC-IM was performed with 100 bootstrap replicates using the options -mu 1.82e-8 -b 1e-8,1e-2. The values of b1 and b2 followed Wang et al. (2022).

Gene flow to the Tibetan population was estimated by the value of *f*_4_(Outgroup, Central or West Asia; East Asia, other groups). We used the American black bear (American_black_bear; Kumar et al. 2017) and the Asiatic black bear (Asiatic_black_bear; Kumar et al. 2017) as outgroups and the European and North American populations as the comparison groups.

## Data and resource availability

Raw sequence reads and mitochondrial DNA sequences are submitted in the DDBJ BioProject database (BioProject ID: PRJDB17730).

## Acknowledgments

We thank Mallory Eckstut, PhD, from Edanz (https://jp.edanz.com/ac) for editing a draft of this manuscript. This work was supported by the Japan Society of the Promotion of Science for Scientific Research (KAKENHI grant nos. 18H05508 and 22KJ0055), the Sasakawa Scientific Research Grant from The Japan Science Society (grant no. 2021-5022), the ZIN project no. 125012800908-0, and the establishment of university fellowships towards the creation of science technology innovation (grant no. JPMJFS2101).

